# Late greenhouse gas mitigation has heterogeneous effects on European caddisfly diversity patterns

**DOI:** 10.1101/411462

**Authors:** Stephanie R Januchowski-Hartley, Christine Lauzeral, Astrid Schmidt-Kloiber, Wolfram Graf, Sebastien Brosse

## Abstract

Little remains known about how the timing of mitigation of current greenhouse gas emissions will influence freshwater biodiversity patterns. Using three general circulation models, we evaluate the response of 260 broad-ranging European caddisfly species to climate conditions in 2080 under two scenarios: business as usual (A2A) and mitigation (A1B). If implemented effectively, recent government commitments established under COP21, to mitigate current greenhouse gas emissions, would result in future climatic conditions similar to the mitigation scenario we explored. Under the Cgcm circulation model, which we found to be the most conservative model, suitable environmental conditions were predicted to shift 3° more to the east under the mitigation scenario compared to business as usual. The majority of broad-ranging European caddisfly species will benefit from mitigation, but 5 to 15% of species that we evaluated will be bigger losers under the mitigation scenario compared to business as usual. Under the mitigation scenario, caddisfly species that will retain less of their current range and experience lower predicted range expansion are those that currently have relatively limited distributions. Continental-scale assessments such as the ones that we present are needed to identify species at greatest risk of range loss under changing climatic conditions.

## INTRODUCTION

A growing number of studies are evaluating how alternative scenarios could influence Earth’s biodiversity under future climate change (McMahon et al. 2011; Garcia et al. 2014; Warren et al. 2018). Series of scenarios have been developed to represent how political decisions influence greenhouse gas emissions and are used to evaluate the subsequent magnitude of policy influences on future climate conditions. Plausible alternative scenarios to business as usual have also been developed to represent the potential benefits gained from mitigating greenhouse gas emissions (Nakicenovic et al. 2000). Immediate and future policy-based actions to reduce greenhouse gas emissions could mitigate the strength of climatic change over the next several decades and reduce biodiversity losses (Nakicenovic 2000).

National level commitments, established ahead of the 21st Conference of the Parties (COP21), aimed to mitigate greenhouse gas emissions through 2025 or 2030 (UNFCC 2015). These commitments are predicted to result in a 3°C increase in surface temperature and climate conditions similar to those depicted under IPCC’s A1B scenario by the end of the century (UNFCC 2015). There remains a need to better understand the influence of such mitigation measures on global and regional biodiversity patterns and processes. Climatic change is also likely to have varied consequences on biodiversity patterns depending on the region considered, and interactions between temperature, precipitation and species-specific tolerances are likely to influence the magnitude and velocity of change in species’ distributions (VanDerWal et al. 2013).

The impact of climatic change on freshwater biodiversity patterns also remains poorly understood (Balint et al. 2011; Domisch et al. 2012). Comte et al. (2013), demonstrated that most of our knowledge about the impact of climate change focused on at least one salmonid species, and that there is a general lack of studies on climate-change effects on threatened species. The situation is similar for freshwater invertebrates, and despite a growing number of studies (Domisch et al. 2012; Simaika et al. 2013; Warren et al. 2018), a broader understanding of potential climate change impacts on this diverse group of species is needed. Literature reviews have been used to evaluate the sensitivity of Europe’s caddisfly species to changing climate (Hering et al. 2009), but to our knowledge only Domisch et al. (2012) have quantified the influence of changing climate on habitat suitability for aquatic macroinvertebrate in Europe.

We explored the potential benefits of mitigating business as usual greenhouse gas emissions for European freshwater biodiversity; focusing on a group of well sampled and broader ranging European caddisfly species. Caddisflies (Trichoptera) constitute a group of interest when it comes to assessing climate change impacts on freshwater biodiversity because they are diverse, and generally broad-ranging, with more than 1700 species in Europe (Graff et al. 2008). We considered current climate, and potential future climate scenarios for 2080 using IPCC scenarios A2A and A1B. We chose these two scenarios because one predicts business as usual emissions (A2A) and the other a leveling off in emissions by 2050 because of mitigation efforts (A1B). We focused our analysis on 260 well-sampled, and relatively broad-ranging, European caddisfly species, and used Iterative Ensemble Models (Lauzeral et al. 2012, 2015) to evaluate how temperature and precipitation changes under these two scenarios and three general circulation models (Cgcm, Hadcm and CSIRO) could modify individual species’ current distributions as well as European-wide species diversity patterns by the end of the 21^st^ century. It is predicted that wide-ranging species will extend their range and that more specialized, range-restricted species will see declines in suitable range areas under future climate conditions (Hering et al. 2009; Domisch et al. 2012). With this in mind, we anticipated that the different climate scenarios we explored would result in varied combinations of both *winners* and *losers* and generate contrasted changes in caddisfly species richness across different areas of Europe.

## METHODS

### Species occurrence data

We extracted caddisfly species occurrence records from a European-wide database (Schmidt-Kloiber et al. 2017). To our knowledge this database is the most detailed and comprehensive database for European Trichoptera. Our assessment started with 322 caddisfly species which had more than 100 records, and a total of 395,513 records in the database. We removed species living in ponds or wetlands from the dataset because air temperature is a poor proxy for the influence of temperature on species dependent on these deeper water habitats (Caissie 2006). Further, only the species with more than 100 occurrence records in the database were considered in our subsequent analysis to ensure more reliable predictions. We also removed individual species occurrence records from before the year 1950, and only retained records up to the year 2000, and did this to ensure that records aligned with the time period of current climatic data considered (1950-2000). We also ensured that individual records retained for modelling had an accuracy of at least 1 km to reduce spatial error.

Our final database contained 260 caddisfly species, whose current distribution areas varied from 3 to 42% of Europe’s total area (mean = 2.4 ± 0.8 million km^2^ SD; range size = 0.3 – 4.2 million km^2^ SD). The 260 modeled ‘current’ distribution ranges also fit in each of the species’ known distributions in European ecoregions; validated by two Trichoptera experts (A. Schmidt Kloiber and W. Graf).

### Climate variables

We accessed global-scale spatial climate data for both current (1950-2000) and future (2080), from WorldClim (http://www.worldclim.org). All spatial climate data were 30 arc-seconds, approximately 1 km x 1 km, spatial resolution. Based on current conditions, we considered only those ecologically relevant climatic variables and removed correlated variables, based on Pearson’s correlation coefficients. When two climatic variables were strongly correlated (r>0.7), we retained the most ecologically relevant variable, resulting in six climatic variables included for all subsequent species distribution modelling: 1) temperature seasonality; 2) maximum temperature of the warmest month; 3) minimum temperature of coldest month; 4) precipitation of wettest month; 5) precipitation of driest month and 6) precipitation seasonality. We assumed air temperature as a substitute for water temperatures, because European-wide data on projected changes in water temperature are not available. Further, caddisflies depend on both aquatic (larval) and terrestrial (adult) environments, and the potential for caddisfly sensitivity to changes in temperature have been previously demonstrated by Hering et al. (2009). Moreover, using air temperature as a substitute for water temperature is generally acceptable for large scale studies that cover a certain extent of climate, because air and water temperature in streams and rivers are strongly positively correlated (Caissie et al. 2006). For 2080, we considered these six climatic variables under A1B and A2A scenarios of anthropogenic activity from the 4^th^ Assessment Report of the Intergovernmental Panel on Climate Change (IPCC 2007), and Cgcm (Canadian Centre for Climate Modelling and Analysis), Hadcm (Hadley Centre for Climate Prediction and Research’s General Circulation Model) and CSIRO (Commonwealth Scientific and Industrial Research Organization) GCMs. The three GCMs we selected have been previously used to evaluate the impact of climate change on freshwater organisms in Europe (Domisch et al. 2012; Buisson et al. 2009). We refrained from averaging across GCMs because the goal of our study was to demonstrate variability between models, and averaging across GCMs can smooth patterns and limit our ability to fully assess alternative scenario influences on climate suitability, and ultimately on species patterns.

### Species distribution models

We modelled current and future distributions for 260 caddisfly species using an ensemble modeling framework developed by Lauzeral et al. (2015). Ensemble models are known to be more efficient than single models for predicting species distributions (Marmion et al. 2009), but they need reliable presence and absence data (Lobo et al. 2010). Presence-only models, such as Maxent provide an alternative to the lack of reliable absences (Phillips et al. 2006), but such models are known to overestimate the range of species (Yackulic et al. 2013; Ward et al. 2009). Iterative Ensemble Models (IEM) offer a way to deal with uncertain absences in ensemble models and have been shown to provide reliable predictions of species distributions (Lauzeral et al. 2012). IEM is an improvement of the ensemble models that simultaneously apply a wide range of statistical methods to produce a consensual response that synthesizes individual model outputs. The iterative step of the IEM enhances models reliability by correcting for incompleteness in species distribution databases (Lauzeral et al. 2012). We determined that IEMs were well suited for our data, where false absences (the species has not been detected, but is present) are likely to be present (Lobo et al. 2010). Indeed, despite more than a century of intensive surveys carried out across Europe (Schmidt-Kloiber et al. 2017), the absence of a given caddisfly species in a European region remains uncertain.

Although criticized for not incorporating ecological processes (Evans et al. 2012), IEMs are considered the most efficient method for predicting species distributions when species’ ecological traits are poorly understood or not available (Araújo et al. 2007). Our IEM used six predictive modelling methods belonging to three commonly used correlative species distribution modelling techniques. We used two regression techniques: generalized linear models (GLM) and generalized additive models (GAM); two machine learning techniques: random forest (RF) and generalized boosted regression models (GBM); and two classification techniques, linear discriminant analysis (LDA) classification and regression trees (CART). Raw variables were used without prior transformation in all models except for GLM and LDA models where variables were squared to deal with nonlinearity, and in the GAM model, where variables were spline transformed (df = 4). We generated 1000 trees in our GBM models and 300 trees in our RF models, and for both of these modelling methods, the number of predictors randomly selected at each node was the square root of the total number of climate variables (n = 6).

The six model outputs from IEM were averaged to provide a per-pixel relative suitability for each species, which was then converted into presence or absence by maximizing the True Skill Statistic (TSS). The calibration data set was randomly selected as 70% of the data matrix. This process was repeated 10 times to measure the sensitivity of our predictions to the calibration dataset, giving rise to 10 presence-absence values per 1 km^2^ pixel. The species was considered as present if predicted in at least 5 out of the 10 repeats. Model quality was quantified using TSS, accounting for model sensitivity and specificity. All statistical analyses and modelling were carried out in R Statistical Software Version 3.1 (http://www.R-project.org/).

Our models predicted current and potential future range distributions for 260 European caddisfly species. Using these predictions, we represented future (2080) species ranges considering both no dispersal and dispersal scenarios for each GCM. Under no dispersal scenarios, species ranges were constrained to their current distribution ranges, and under dispersal scenarios predicted species ranges extended outside their existing distribution range.

## RESULTS

Our models showed good performance for each of the 260 caddisfly species (TSS > 0.6), with a mean TSS = 0.83 (± 0.06 SD) and low variability in model performance across species. Based on the 260 caddisfly species considered in our analysis, we found that species richness peaks in central Europe (Fig. 1a). Under a non-dispersal scenario, species richness would decline throughout Europe regardless of the scenario (Fig. 1b) or the circulation model considered (Fig. 1b, S1b and S2b). In addition, under a non-dispersal scenario, mitigation primarily benefits species in areas of Central and Eastern Europe, whereas under mitigation, Southern Europe (e.g. areas of Italy and Greece; Fig. 1b) loses more species.

**Figure 1.**
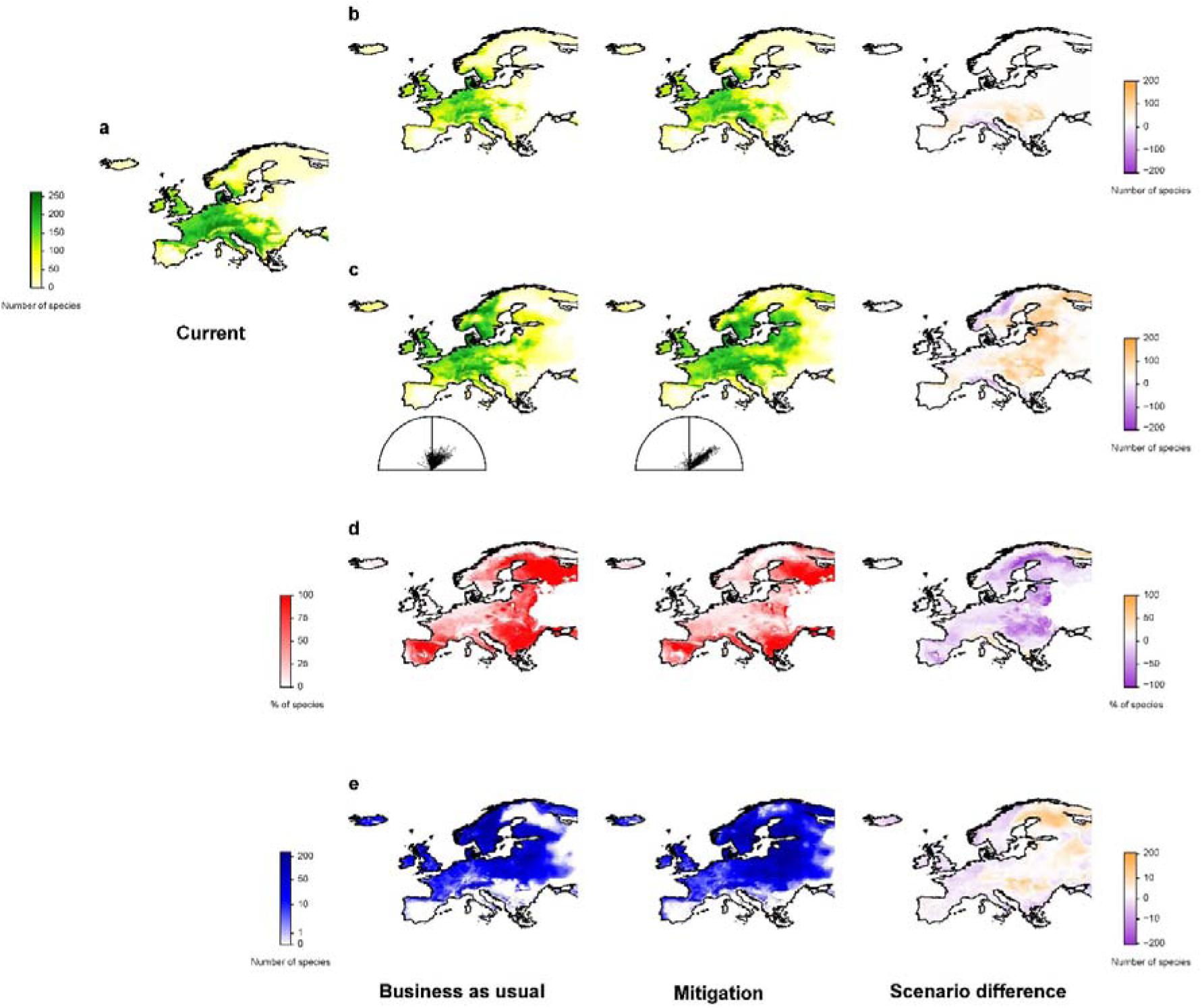
Current and predicted biodiversity patterns for 260 European Trichoptera species. Biodiversity patterns for: (a) current species richness and using four metrics to assess future patterns: (b) species richness under no dispersal, (c) species richness under dispersal, (d) the percentage of species lost per pixel and (e) the number of species gained per pixel compared to current distributions. The four metrics are depicted based on business as usual (A2A) and mitigation (A1B) scenarios, using Cgcm General Circulation Model. The difference in the number or percentage of species per pixel between mitigation and business as usual scenarios is on the right panel for b, c and d. Higher values under the mitigation scenario are positive values. The half circles represent the strength and directionality of movement in the centroid of each species’ distribution under the business as usual and mitigation scenarios, respectively. All images were created using R Statistical Software Version 3.1 (http://www.R-project.org).

Similar to the non-dispersal scenario, when allowing for species’ dispersal, areas of Southern Europe (Italy and Greece; Fig. 1c) lose more species under the mitigation scenario. Allowing species dispersal results in species richness shifting in both a north and east direction by 2080, regardless of the circulation model considered (Fig. 1c, S1c and S2c). Using Cgcm GCM, which provides the most conservative shifts in species distributions, the northward shifts in the centroid of caddisfly species’ distributions are 4.87±1.03°SD under business as usual and 4.93±1.34° SD under the mitigation scenario, with no significant difference between scenarios (t-test, p>0.23). In contrast, the magnitude of eastern shift in species richness significantly differs between scenarios (t-test, p <0.01), and surprisingly, the centroid of richness shifts three degrees further to the east under the mitigation scenario (4.47±2.56°SD) compared to business as usual (1.33±2.24°SD) (Fig. 1c).

The Cgcm GCM predicts increased suitability, with caddisfly species richness increasing across 64% of the European landscape under the mitigation scenario compared to under business as usual (Fig. 1c). Our predictions also show that most of the European landscape (55% of total area) is predicted to experience higher species loss under business as usual (Fig. 1d). However, under the mitigation scenario, 16% of Europe has more pronounced species loss and 40% of Europe experiences similar loss under both mitigation and business as usual (Fig. 1d). Areas predicted to experience higher species loss under mitigation are in northern Europe as well as parts of Italy and Greece (Fig. 1d). Under mitigation, Northern and Eastern Europe as well as some parts of Spain and Portugal gain higher numbers of species than under business as usual (Fig. 1e). We found similar changes in geographical patterns across Europe under the mitigation scenario for the two other GCMs used (Fig. S1d,e and S2d,e).

We further explored which climatic variables explain predicted differences in species richness patterns between the two future scenarios. Under Cgcm GCM, the difference between the two scenarios in predicted loss or gain of species (measured per pixel) is mainly due to two climate variables (Fig. 2 and S3). Predicted differences in species loss are a consequence of higher maximum temperature of the warmest month predicted across southern Europe under the mitigation scenario (Fig. 2a). Predicted differences in species-gain (per pixel) are a consequence of higher precipitation predicted in the driest month under mitigation (Fig. 2b).

**Figure 2.**
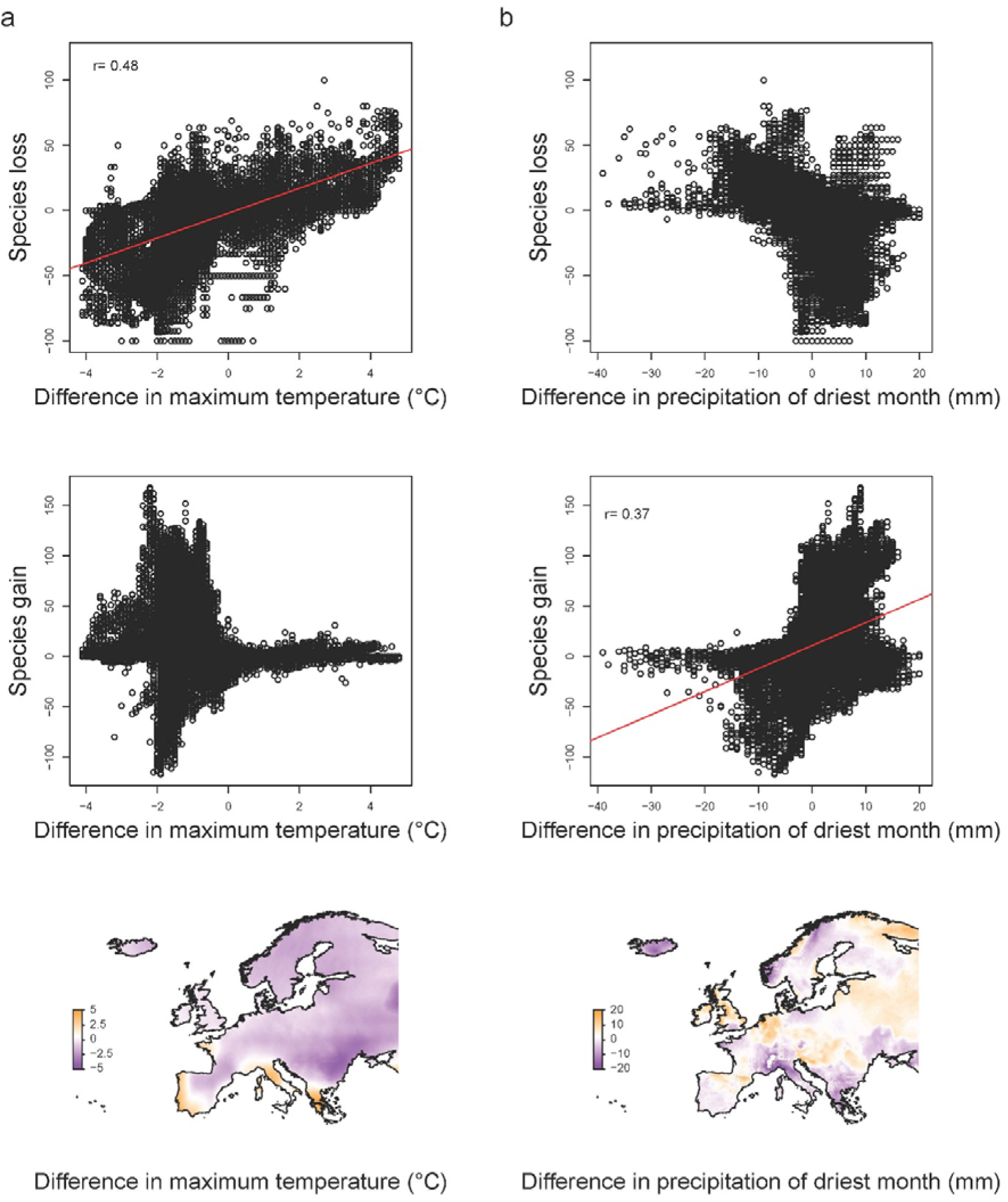
Relationships between future biodiversity patterns and climate variables. Scatterplots of the relationship between the differences in number of species predicted to be lost or gained (per pixel) and difference in (a) maximum temperature and (b) precipitation of driest month under mitigation (A1B) compared to business as usual (A2A) scenario, using Cgcm general circulation model. The regression line is only shown when r > 0.30. The Pearson’s correlation coefficient is given on the top left of each scatterplot. The map insets depict the geographical differences in (a) maximum temperature and (b) precipitation of the driest month between mitigation and business as usual scenarios, where higher values under the mitigation scenario are positive values and depicted in shades of orange. All images were created using R Statistical Software Version 3.1 (http://www.R-project.org).

At the individual species level, species show heterogeneous responses in distribution according to the GCM considered. On average, species retain 41 to 71% of their current distribution and tend to expand beyond their current distribution by 42 to 97% (Fig. 3, S4 and S5). The effect of mitigating greenhouse gas emissions is also predicted to have heterogeneous effects across GCMs, with Cgcm maintaining highest proportion of species’ current distributions (Fig. 3, S4 and S5). On average, under Cgcm, species retain 5% more of their current distribution under a mitigation compared to business as usual scenario, but also expand their distribution by the year 2080 (23% of their current range on average) under the mitigation scenario (Fig. 3).

**Figure 3.**
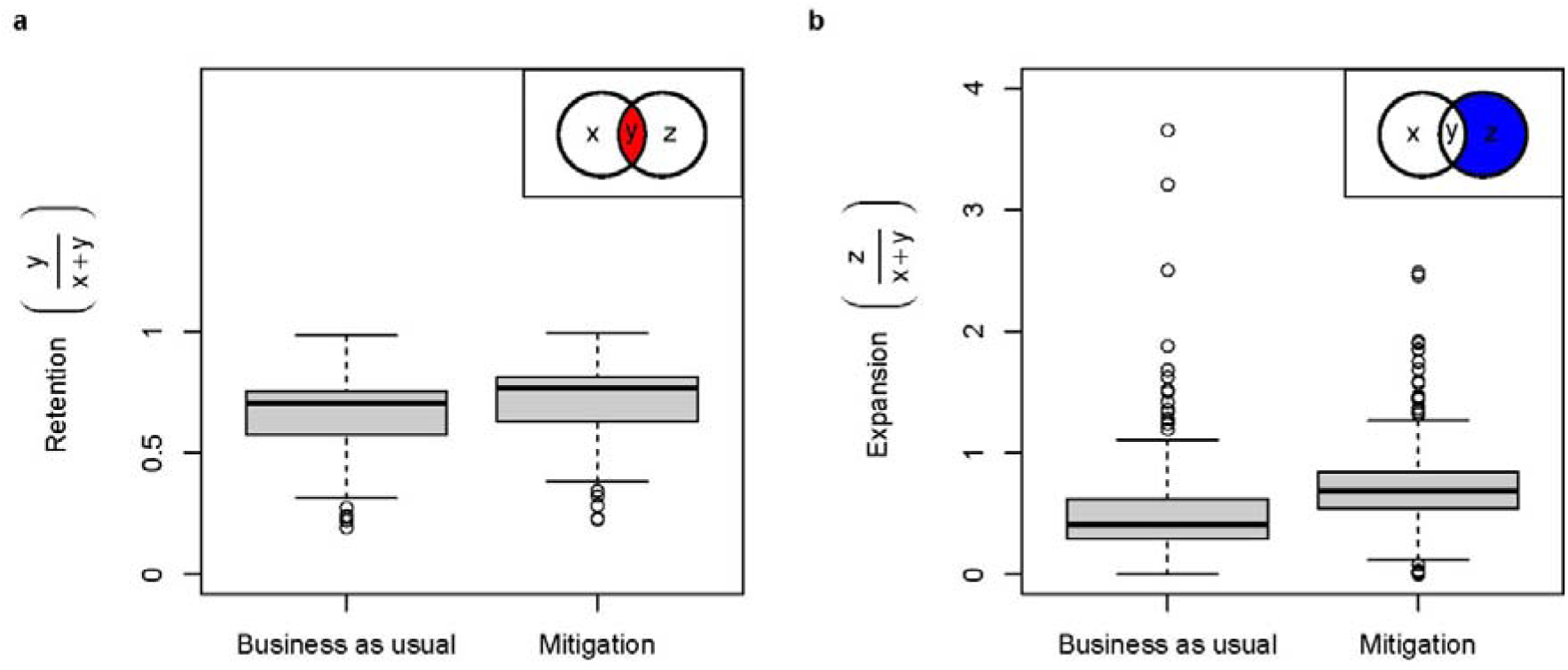
Retention and expansion of species’ distributions under future scenarios. Boxplots represent the retention and expansion of species’ distributions between current and business as usual (A2A) and mitigation (A1B) scenarios, using Cgcm general circulation model. The inset of each boxplot illustrates hypothetical current (left circle) and future (right circle) distributions of a species, where (x) is the current area that could be lost, (y) is the current area retained in future and (z) is the new area predicted in future. Retention is the proportion of a species’ current geographical distribution area which persists under future climate conditions. Expansion is the predicted distribution area outside of a species’ current distribution area divided by current distribution area. An expansion value greater than one means a species is predicted to colonize a larger area than its current distribution area. All images were created using R Statistical Software Version 3.1 (http://www.R-project.org).

Roughly 20% of species (50 species) in our study are predicted to be losers, either retaining less of their current distribution (37 species) or expanding less into new areas (28 species) under the mitigation scenario compared to business as usual (Fig. 4). Species with relatively limited distributions in mountainous areas, parts of the Mediterranean and extreme north Europe, are predicted to be at greater risk of distribution loss under mitigation, using Cgcm and CSIRO GCMs (Fig. 4 and S6). For instance, under mitigation, the majority of predicted ‘losers’ tend to be species that currently have relatively limited distributions (18% of total European area based on the 50 ‘loser’ species; Fig. 4). Hadcm GCM predicts reduced benefit to species from mitigation, and losers are more widely distributed across Europe (Fig. S7).

**Figure 4.**
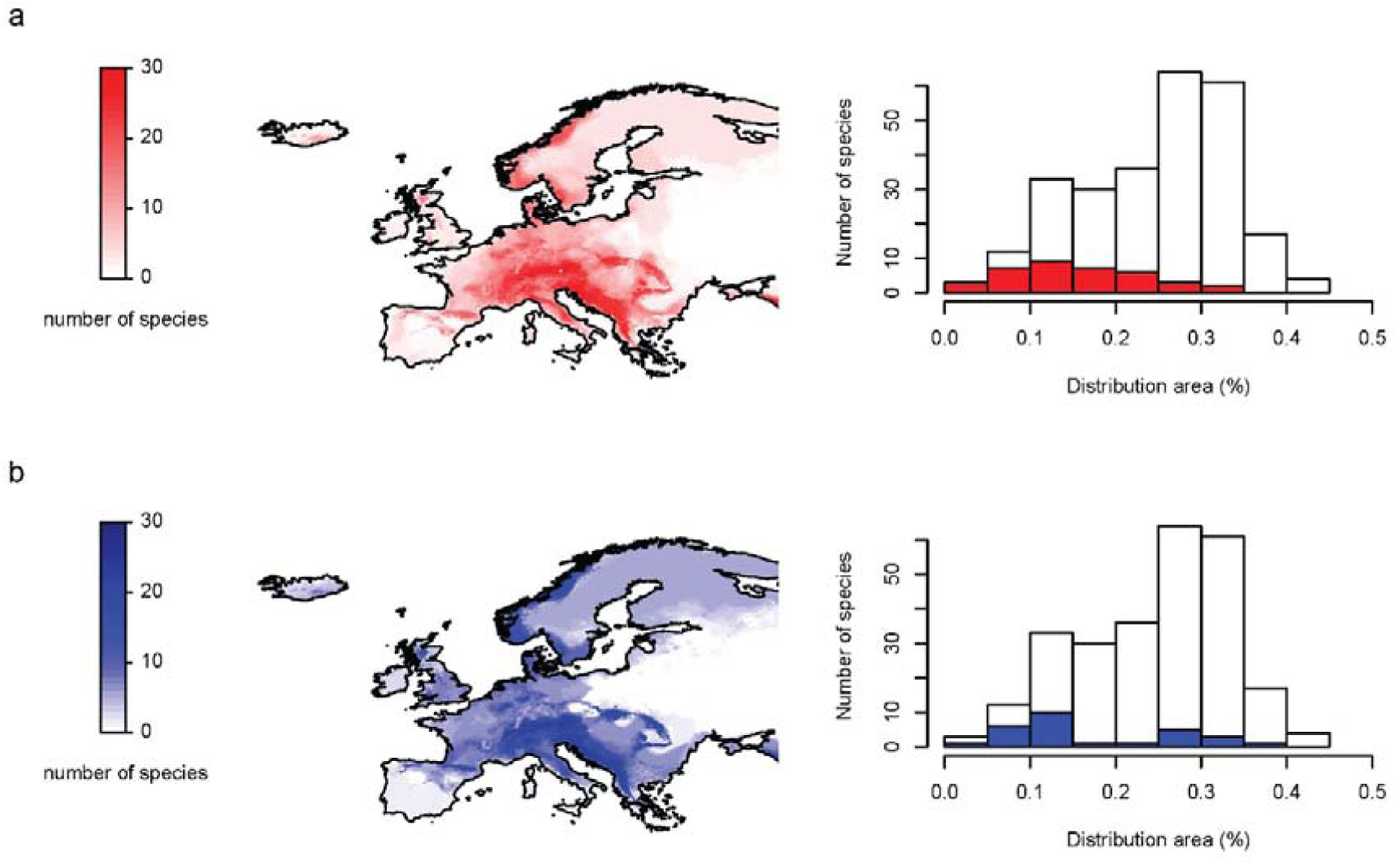
Metrics of species’ that will not benefit from mitigation. Maps represent the current richness (per pixel) of those caddisfly species (n = 50) which are predicted to be bigger losers under mitigation (A1B) than business as usual (A2A), using the Cgcm General Circulation Model. Losers are either species predicted to have (a) less retention of their current distribution area or (b) less expansion of distribution area under mitigation compared to business as usual. Histograms of current distribution area occupied by 260 caddisfly species (white bars) and species which are predicted to have (a) less retention of their current distribution area (in red) or (b) less expansion of distribution area (in blue) under mitigation compared to business as usual. Distribution area is represented as the proportional area of Europe that a species currently occupies. All images were created using R Statistical Software Version 3.1 (http://www.R-project.org).

## DISCUSSION

Our findings suggest that even late mitigation of greenhouse gas emissions, as depicted under Cgcm GCM, will maximize retention of current European distribution areas for most broader ranging caddisfly species compared to maintaining business as usual. However, we also found that a mitigation scenario will have heterogeneous effects on species distributions depending both on the species considered and global circulation conditions. The ecological consequences of heterogeneous effects on species distributions remain poorly understood, and to our knowledge no studies have evaluated the potential implications of possible changes in species composition on food-web dynamics or the maintenance of important ecological processes in Europe’s freshwater ecosystems. This remains an open area for research and would provide improved understanding about how climate change could influence freshwater ecological processes at regional scales.

Mitigation efforts, as depicted under A1B scenario and Cgcm GCM, are predicted to put 14% of the caddisfly species we considered in our study at greater risk of losing distributional area than under business as usual. Our results suggest that mitigating climate change by 2050 will not linearly lower changes or impacts to caddisfly species – some of the broader ranging species considered in our analysis stand to lose regardless of these efforts. Indeed, even though climatic conditions will be globally improved under mitigation, in a few places, climate change is predicted to be more pronounced under mitigation than under business as usual. For instance, we found that under the mitigation scenario we considered that temperature is predicted to reach higher values in Western and Southern Spain, Italy and Greece. Despite heterogeneities in our model responses according to the GCM considered, all the models showed that species currently inhabiting Southern France, Italy and the Balkans will benefit the least from efforts to mitigation greenhouse gasses by 2050. These areas, Southern France, Italy, and the Balkans also host high caddisfly species endemicity – species that Hering et al. (2009) suggest will have limited ability to adapt to changing climate.

When considering both a no-dispersal and a dispersal scenario we found a decline in species richness in Southern Europe. However, we found that if species were able to freely disperse then species richness would increase in both Eastern and Northern Europe by 2080. Caddisflies are relatively poor dispersers compared to other flying macroinvertebrates like dragonflies, but large ranging caddisfly species, like those considered in our study, are known to be better dispersers compared to species with more restricted ranges (Hering et al. 2009). We were unable to account for individual species dispersal abilities because this information is known for so few species. It is possible that explicit consideration of species’ dispersal abilities, as opposed to unlimited dispersal, would restrict the potential expansion of species into new regions and identify even greater losses for species. In turn, our dispersal scenarios offer a conservative view, and are likely to exceed most species actual dispersal abilities. Despite this limitation it is important to evaluate scenarios that consider potential dispersal even though specific dispersal abilities remain poorly understood (Chen et al. 2011; Heino et al. 2009). In addition to our limited ability to account for species’ dispersal, we were not able to account for other human disturbances or hydrological conditions into the future. As noted above this means that our predictions likely offer an optimistic view of how caddisfly species distributions in Europe are likely to be affected under climate change and overcoming the limitations of our study would likely identify additional negative impacts of climate change on habitat availability and possibly even greater predicted loss of species.

Our modelling approach also required us to focus on relatively broad-ranging, data rich, species, meaning our results could overlook additional species loss from mountain tops or small localized areas where species with relatively restricted distributions occur. Therefore, overall patterns observed in our study are likely to be further emphasized by including species with narrower distributions that are also considered to be more sensitive to climate change, such as those inhabiting mountains or mountainous areas. Given the high likelihood of these climatic conditions in future, proactive strategies are needed to identify species that will potentially not benefit from climate change mitigation efforts and to identify strategies (e.g., species translocations; mitigation of other human-disturbances) to mitigate impacts. There could be great benefit in more explicitly examining both no dispersal and dispersal scenarios in relation to species sensitivity to climate change – characteristics outlined by Hering et al. (2009). For example, Hering et al. (2009) demonstrate the status quo of species vulnerability to climate change, but coupling data generated from their research with the models generated here, would allow for a more dynamic and proactive approach. Coupling these methods could help us to determine how changes in species distributions further influences their sensitivity to climate change, and to also identify regions where sensitive species could be supported in future.

## ACKNOWLEDGEMENTS

This study was supported by the BioFresh European project (FP7-ENV-2008; contract number 226874). We thank F. Januchowski-Hartley and S. Vitecek for feedback on earlier versions of this manuscript.

